# Critical Residues of Gβγ for the interaction with the SNARE Complex

**DOI:** 10.1101/2020.04.29.069187

**Authors:** Benjamin K. Mueller, Ali I Kaya, Zack Zurawski, Yun Young Yim, Jens Meiler, Heidi E. Hamm

**Author notes:** NE-CAT, Department of Chemistry and Chemical Biology, Cornell University, Argonne National Laboratory, Argonne, IL 60439, USA. Department of Anatomy and Cell Biology, University of Illinois at Chicago, Chicago, IL 60612, USA. Nash Department of Neuroscience, Icahn School of Medicine at Mount Sinai, New York, NY 10029, USA. Correspondence: Heidi E. Hamm.

## Abstract

The mechanisms and regulation of neurotransmitter release is a complex process involving many co-factors and proteins. One critical interaction is the regulation of exocytosis when G-protein βγ (Gβγ) dimers bind to the soluble N-ethylmaleimide-sensitive factor attachment protein receptor (SNARE) protein complex. The complex is comprised of N-ethylmaleimide-sensitive factor attachment protein-25 (SNAP-25), syntaxin 1A, and synaptobrevin. Herein we probe across the entire family of human Gβ and Gγ proteins for residues critical for the interaction with SNARE, by systematically screening peptide sequences for their ability to bind to tSNARE. The coiled-coil region of Gβγ showed high affinity to tSNARE, along with the propeller region of Gβ on the opposite side from the coiled-coil region. Peptides based on Gβ_1_γ_2_, shown to have high affinity to SNARE, tSNARE were screened further by alanine scanning to probe for residues critical for binding to tSNARE. Full length Gβ_1_γ_2_ and SNARE were docked computationally using Rosetta, to examine the experimentally determined binding sites. Docking converged on two possible sites of interaction using two distinct regions of both Gβ_1_γ_2_ and SNARE.

## Introduction

Exocytosis is a complex, regulated process involving the exocytotic machinery proteins, synaptic proteins that play roles in docking and priming the vesicle, ion channels, calcium sensors, and pre-synaptic inhibitory G-protein coupled receptors (GPCRs). Despite numerous attempts to understand the synaptic molecular composition and function, the microarchitecture of exocytosis with various modulators and synaptic proteins at each step has still remained unclear.

Gβγ subunits mediate fast, membrane-delimited effects on transmission, through both postsynaptic activation of GIRK channels (Logothetis et al., 1987; Reuveny et al., 1994) and binding voltage-gated calcium channels (VGCC) and modulation of calcium entry (Herlitze, 1996; Ikeda, 1996). At presynaptic terminals, Gβγ is an important regulator of neurotransmission through interactions with calcium channels (Herlitze, 1996; Ikeda, 1996) and via it’s interaction with the SNARE complex which directly inhibits exocytosis (Betke et al., 2012; Blackmer et al., 2005; Blackmer et al., 2001; Gerachshenko et al., 2005). The interaction of Gβγ with the SNARE complex places inhibitory regulation of exocytosis at the final step of exocytotic fusion (Wells et al., 2012b).

There are five different Gβ and twelve different Gγ subunits (Downes and Gautam, 1999; Hildebrandt, 1997; Simon et al., 1991). Gβ subunits are made up of an α-helix of approximately 20 amino acids and 7 blades of four anti--parallel β-strands, a propeller like WD repeat (Clapham and Neer, 1997; Gautam et al., 1998; Smrcka, 2008; Sondek et al., 1996b). Gβ1-4 share up to 90% amino acid sequence identity whereas Gβ_5_ is only 50% identical (Betty et al., 1998; Smrcka, 2008). In contrast, Gγ subunits share only 30-70% sequence identity (Betty et al., 1998; Smrcka, 2008). Made up of two α-helices, the C-terminal α helix interacts with the surface of blade 5 and a small section of the N--- terminal region on Gβ while the N-terminal α-helix forms a coiled-coil interaction with the N-terminal helix of the β subunit (Clapham and Neer, 1997). Gγ subunits can be post-translationally modified at the processed C-terminal cysteine which is carboxymethylated and modified with a farnesyl or geranylgeranyl moiety via a thioether bond. These modifications mediate membrane localization of Gβγ (Clarke, 1992; Cox, 1995). Together, Gβ and Gγ subunits form Gβγ dimers, and once assembled, act as signaling units for GPCRs and modulate synaptic transmission by interacting with a number of effectors such as voltage-gated calcium (VGCC) and inward-rectifying potassium (GIRK) channels, and soluble NSF attachment protein receptors (SNARE) to regulate neurotransmitter release at the synapse (Blackmer et al., 2001; Currie, 2010; Gerachshenko et al., 2005; Herlitze et al., 1996; Huang et al., 1995; Sadja and Reuveny, 2009; Wells et al., 2012a; Yoon et al., 2007).

SNAP25 is the most important Gβγ partner in the interaction between Gβγ and SNARE, a trimer of SNAP25, syntaxin 1A, and synaptobrevin (VAMP) and in the interaction between Gβγ and the tSNARE dimer, SNAP25 with syntaxin 1A (Blackmer et al., 2005; Blackmer et al., 2001; Delaney et al., 2007; Gerachshenko et al., 2005; Photowala et al., 2006; Yoon et al., 2007; Yoon et al., 2008; Zhang et al., 2011; Zhao et al., 2010). Multiple studies have shown the carboxy(C)-terminus of SNAP25 to be important for the exocytotic function and its interaction with Gβγ -botulinum toxin A (BoNT/A) cleaves the C-terminus of SNAP25 reduces Gβγ/SNARE binding, and a 14-amino acid peptide of the C-terminus of SNAP25 inhibits the formation of the Gβγ/SNARE complex (Binz et al., 1994; Blackmer et al., 2005; Gerachshenko et al., 2005; Schiavo et al., 1993; Yoon et al., 2007; Zhao et al., 2010). In Wells *et al, in vitro* binding studies have demonstrated direct Gβγ-SNARE interaction sites at both the vesicle proximal C-termus as well as the other end of the SNARE complex where zipping up begins, as well as in the H_3_ domain of syntaxin (Wells et al., 2012a; Yoon et al., 2007). In addition, Asp99 and Lys102 in the linker region between the two helices of SNAP25 in close proximity to its palmitoylation sites (Cys 85, 88, 90, and 92) may be important for this interaction. Interestingly, Gly63 and Met64 that had significant reduction in binding are buried in the interface with syntaxin 1A and VAMP.

Despite numerous studies probing the sites on the SNARE complex of Gβγ interaction, it is still unclear where on Gβγ the interaction sites are. The Gβγ-effector interaction sites were hypothesized to be localized at the Gα interface (Ford et al., 1998). However, mutagenesis studies revealed the binding regions of effectors within the blades of the Gβ propeller and its N-- terminus coiled--coil, the Gα subunit interface, and regions on Gγ subunit (Clapham and Neer, 1997; Myung et al., 2006; Panchenko et al., 1998; Peng et al., 2003; Smrcka, 2008). For example, GIRK channels were found to interact just outside of the Gα binding site. Thr 86, Thr87, and Gly131 on Gβ_1_ are important for the Gβγ-GIRK interaction (Zhao et al., 2003). The Gα-interacting residues that were shown to be important for Gβγ interaction with GIRK, VGCC, PLCβ, GRK2, and Gα (Ford et al., 1998) were shown to actually be inhibitory for interaction with the SNARE complex and inhibition of exocytosis; removal of those residues and replacement with Ala actually increases their affinity (Blackmer et al., 2005). Mutation of both Lys78 and Trp332 on Gβ_1_γ_2_ to Ala increased the affinity of Gβ_1_γ_2_ for tSNARE by two-fold (Zurawski et al., 2017b). While the interface of Gα subunits likely represents a core site of Gβγ and its effector binding, other regions may also play important roles in facilitating Gβγ mediated downstream signaling.

We have recently reported specificity of Gβγ interaction with α_2a_AR and the SNARE complex based on proteomic studies (Yim et al., 2019) (Yim et al., 2020). Here, we address the structural basis of the specificity of different Gβ and Gγ subunits for the SNARE complex using the method of peptide arrays (Yim et al., 2015a) to further understand the Gβγ-SNARE interaction. The interaction is probed further by computationally docking Gβγ to SNARE, and evaluating the models against the mutagenesis screens.

## Results

To characterize the tSNARE binding sites on G protein β and γ subunits, we used a peptide array assay. Peptides from across the sequences of human Gβ and Gγ protein were synthesized by using Respep SL as described (Yim et al., 2015b). The spots on nylon membrane consisted of 15-mer peptides shifting three amino acids for each successive peptide which cumulatively represents the entire sequence of indicated proteins. **Fig 1A** shows a representative blot and the average data for the interaction between Gβ_1_ and tSNARE. The data revealed 6 major regions of Gβ_1_ that make up the interaction with tSNARE protein. When we tested other Gβ subunits, we found similar binding pattern between Gβ_1-4_ and tSNARE **(Fig 1B)**. Gβ_5_ showed low binding affinity compare to others. This binding data suggests that Gβ_1-4_ has multiple interacting sites with tSNARE protein **(Fig 1C)**.

**Fig 1.**
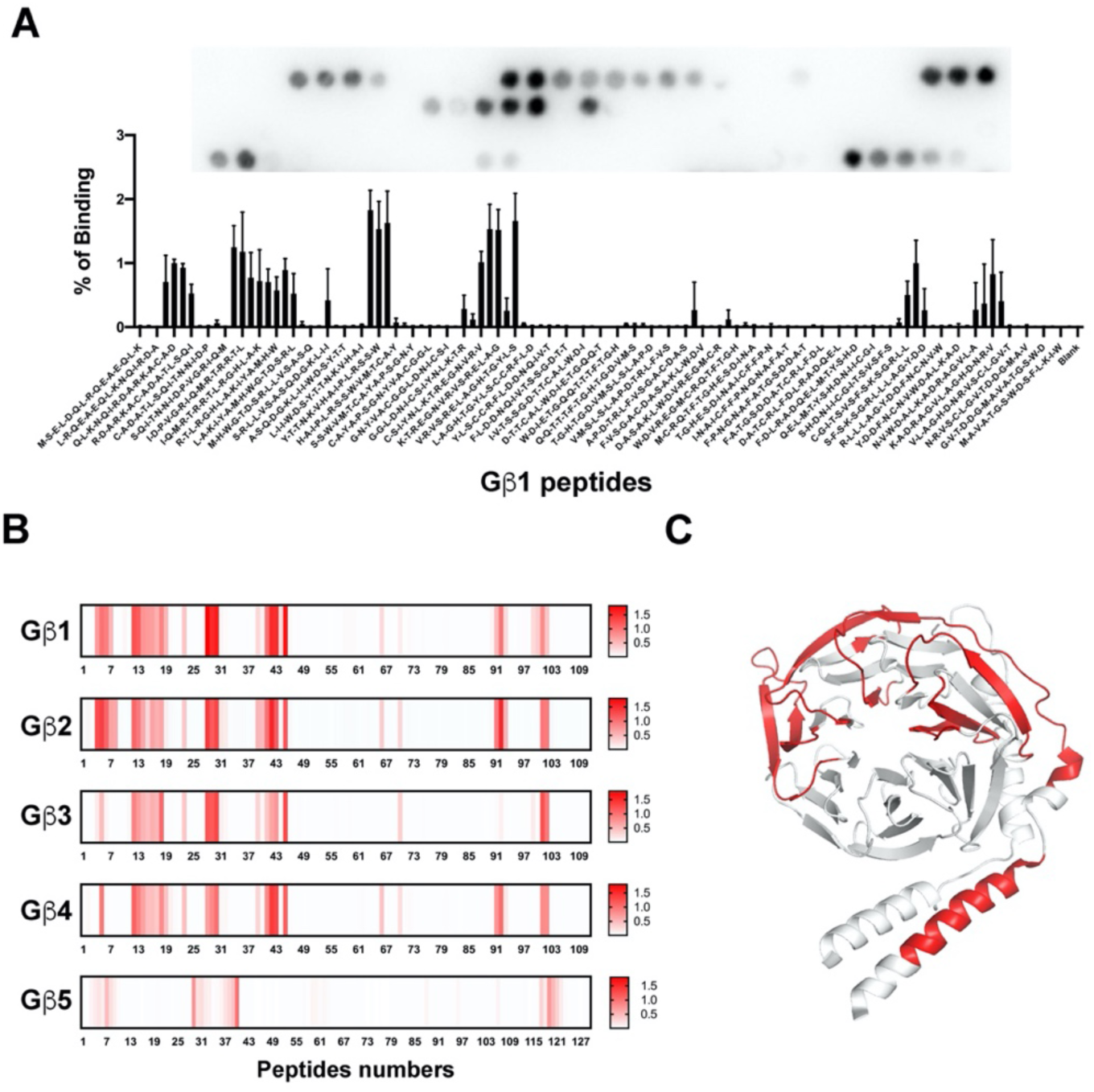
tSNARE binding sites on Gβ subunits. **A)** Representative peptide blot (15-mer) of human Gβ1 and quantified binding of tSNARE to Gβ_1_ peptides. Binding was measured as a percentage of total intensity on the membrane. A 15-mer glycine peptide was used to determine non-specific interactions. Human SNAP25 peptides between amino acids 91-105 and 106-121 were used as positive control for SNAP25 antibody **B)** Heat map using a single gradient to display ranges of binding between tSNARE and Gβ_1-5_ was calculated using GraphPad Prism 8.0.2. **C)** Map of interaction sites for tSNARE to Gβ_1_ are highlighted in red (PDB 1GP2(Wall et al., 1995)).

We used the same approach to detect interacting regions of Gγ with tSNARE. Figure 2A shows binding between Gγ_1_ and Gγ_2_ and tSNARE. The data clearly indicate that the binding is significantly higher with Gγ_2_ peptides than Gγ_1_ **(Fig. 2A)**. It appears that there are two major binding sites between Gγ_2_ and tSNARE, one at the N-terminus and the other at the C-terminus of Gγ_2_ **(Fig. 2B)**. This is consistent with our previous study(Zurawski et al., 2017a). When we investigated the other Gγ subtypes, we observed similar binding patterns as Gγ_2_ **(Fig. 2C)**. This result indicates that Gγ_1_ has lower affinity to tSNARE, and suggests that the other Gγ subtypes have two binding sites at the N- and the C-termini of the protein.

**Fig 2.**
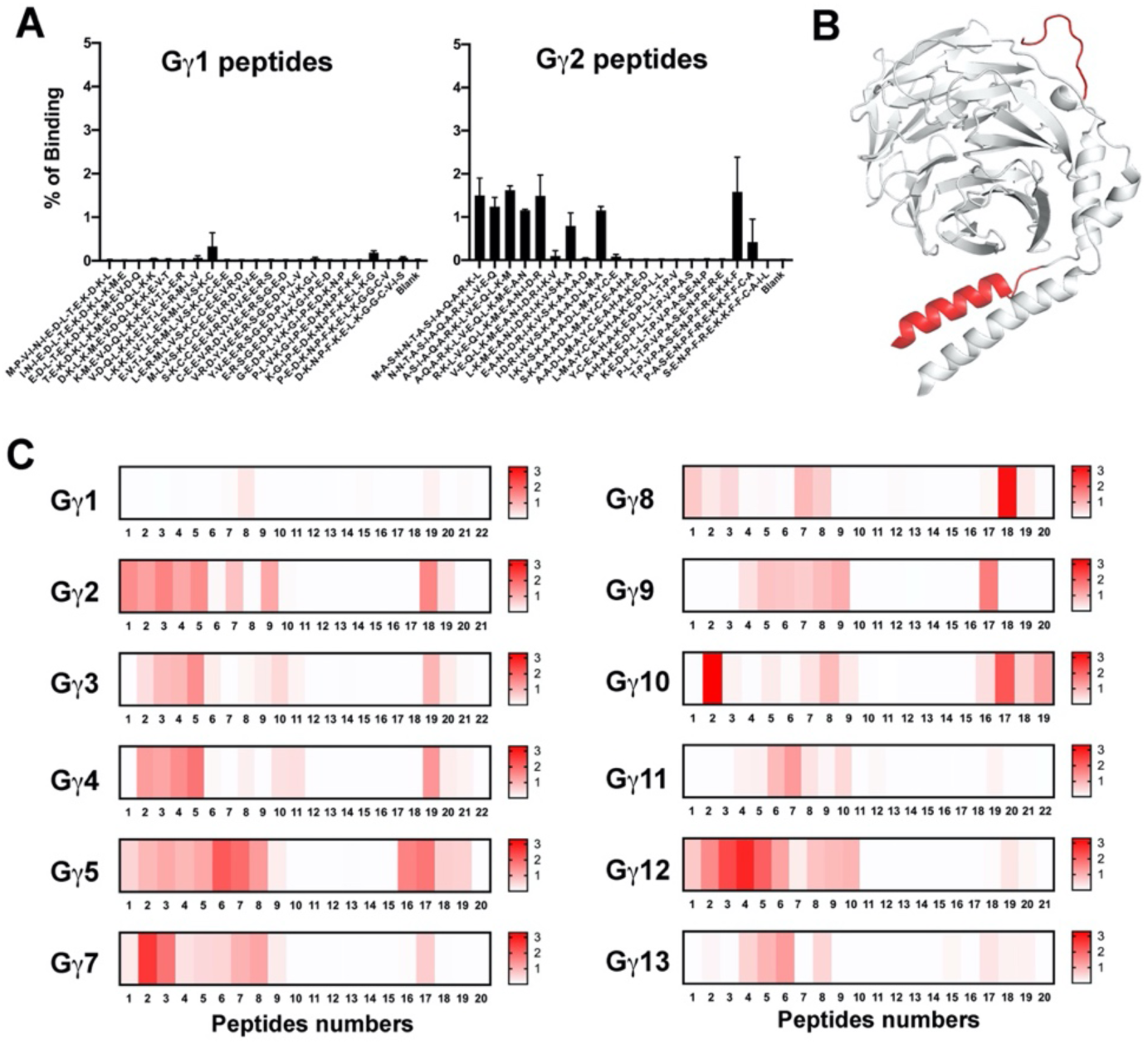
tSNARE binding sites on Gγ subunits. **A)** Representative quantified binding of tSNARE to Gγ_1_ and Gγ_2_ peptides. Binding was measured as a percentage of total intensity on the membrane. A 15-mer glycine peptide was used to determine non-specific interactions. **B)** Map of interaction sites for tSNARE to Gγ_2_ are highlighted in red (PDB 1GP2 (Wall et al., 1995)). **C)** Heat map using a single gradient to display ranges of binding between tSNARE and Gγ_1-13_ was calculated using GraphPad Prism 8.0.2.

On the basis of these results, selected peptides that bound tSNARE in this screen were then subjected to alanine-scanning mutagenesis. For each peptide, the first spot was the wild-type sequence of a given peptide followed by spots with a single alanine mutation. **Fig 3A** shows alanine scanning from Gβ_1_ and Gγ_2_ peptides. The bar graphs demonstrates the average densitometry of all peptides with loss (or gain) of binding in a mutant. The result indicate that positively charged residues play a key role for both Gβ_1_ and Gγ_2_ interaction with tSNARE protein. **Fig 3B** shows the location of the peptides (green) and important residues (blue) for tSNARE interaction.

**Fig 3.**
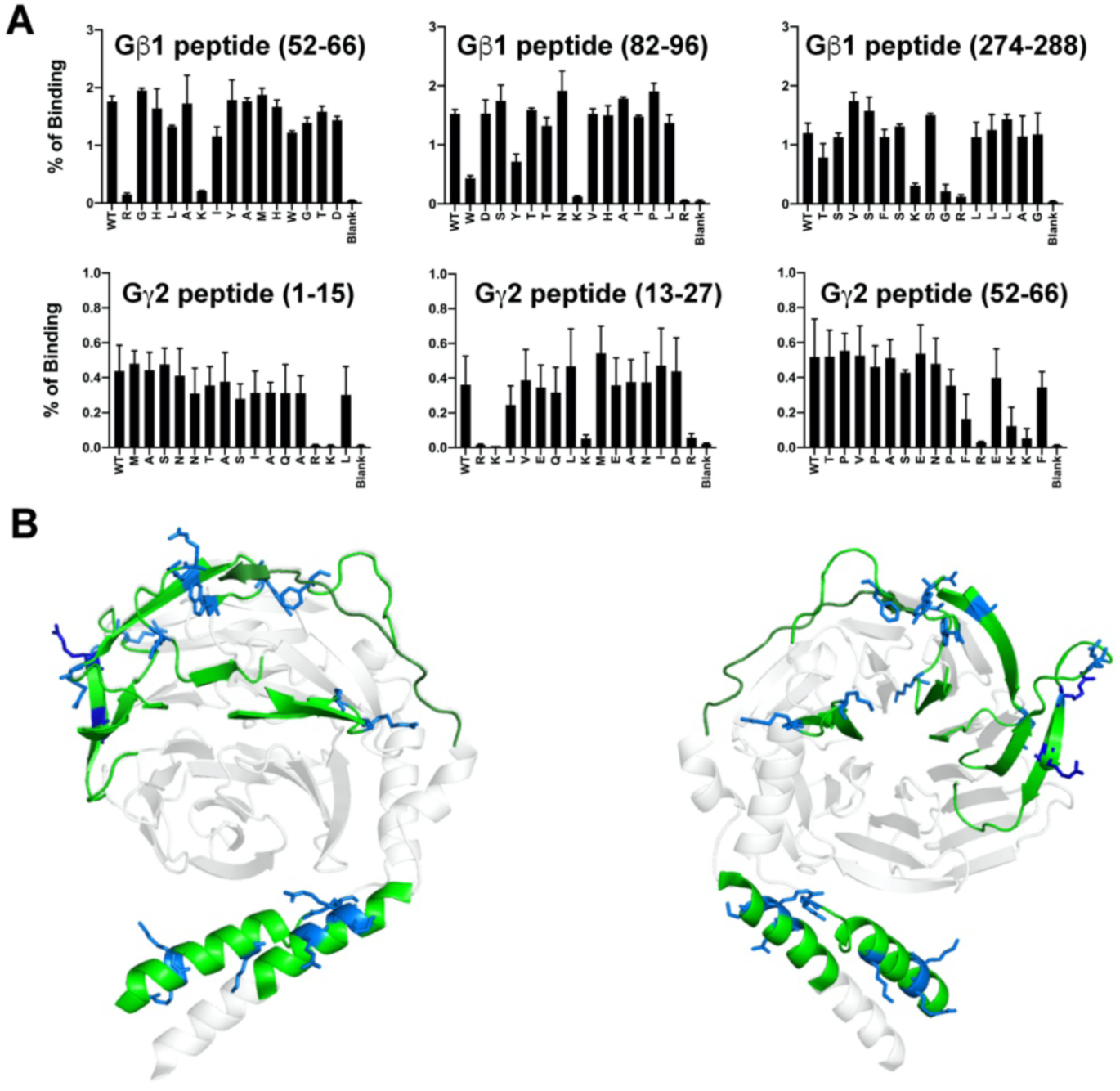
Alanine mutagenesis screening of Gβγ peptides that bind tSNARE. **A)** Selected peptide regions from Gβ_1_ and Gγ_2_ subunits were mutated to alanine. The first bar of each graph contains wild-type peptide. The next 15 bars of each graph to the right are mutant peptides with a single alanine replacement of the residue at position 1, 2, 3. …, 15 for each wild-type peptide. **B)** The peptides are mapped onto 1GP2 structure Gβγ subunit and shown in green. The residues that had significantly reduced binding between Gβγ with tSNARE are shown with blue (stick).

Previous studies showed that the affinity of Gβ_1_γ_1_ is much lower than Gβ_1_γ_2_ for both monomeric SNAP25, t-SNARE, and the ternary SNARE complex. Since the only difference is within the Gγ subunit between these two G protein subtypes, we decided to investigate the affinity of different Gγ subunits to SNAP25. We measured the ability of peptides to disrupt the interaction between full-length Gβ_1_γ_2_ with SNAP25 using the Alphascreen competition-binding assay (Zurawski et al., 2017a). **Fig 4** shows the IC50 differences between Gγ_1_ and Gγ_2_. This result is consistent with previous studies (Zurawski et al., 2017a). We also tested other Gγ subtype peptides (**Table 1**) and the data indicate that Gγ subtypes showing varying affinities for SNAP25. For instance, Gγ_2_ and Gγ_10_ N-terminus and Gγ_8_ C-terminus has higher affinity for SNAP25 compared to other peptides.

**Table 1.**
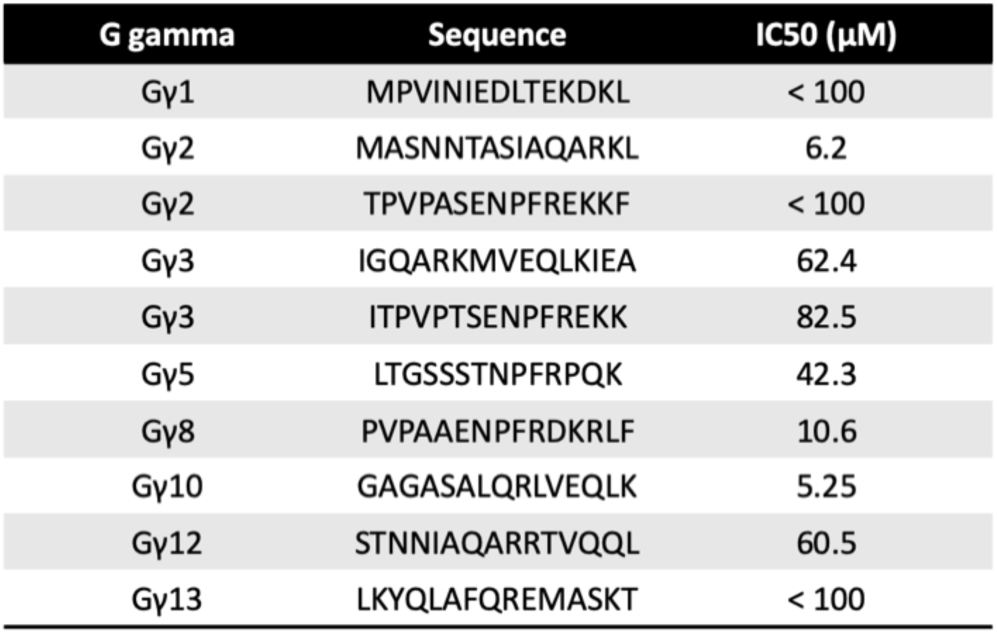
Alphascreen competition-binding assays. Competition assays between biotinylated SNAP25 and Gβ_1_γ_2_ were conducted in the presence of indicated Gγ peptides.

**Fig 4.**
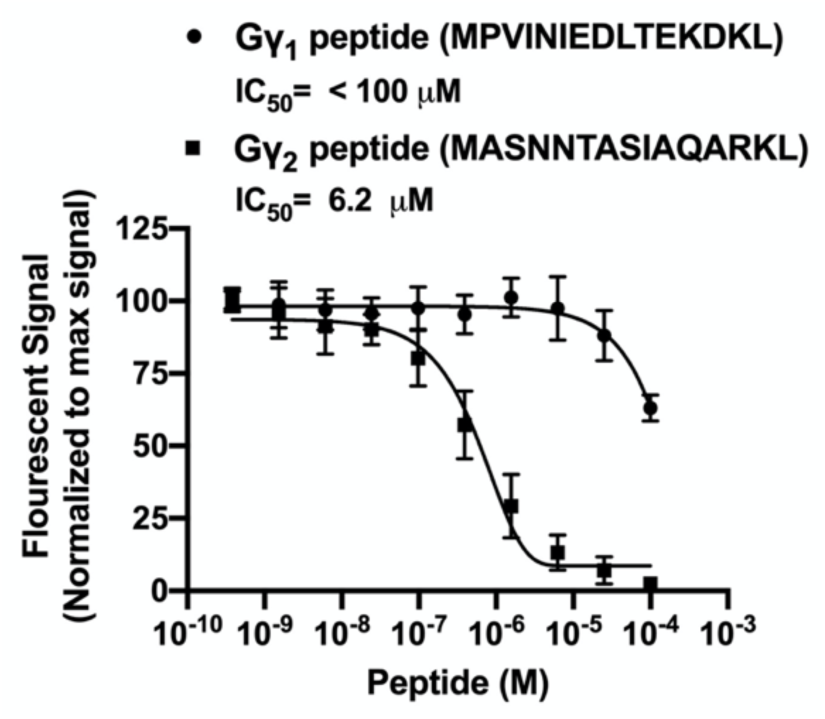
Competition assays of Gβγ-derived peptides to inhibit Gβ_1_γ_2_/SNAP-25 complex formation. Alphascreen competition-binding assays in which 20 nM biotinylated SNAP25 reacts with 180 nM Gβ_1_γ_2_ to produce a luminescent signal were conducted in the presence of peptides corresponding to primary sequences within Gγ1 and Gγ2 peptides at varying concentrations (1.08 nM to 100 µM) dissolved in DMSO. A value of 100% was assigned to the average of all conditions tested containing only DMSO as a positive control. The primary sequence of each peptide is depicted for each condition. Experiments were repeated three times. IC_50_ values and 95% confidence intervals were observed. IC50 of the selected Gγ peptides are shown in **Table 1**.

The interaction between Gβ_1_γ_2_ and SNARE was further probed via computational docking. The structures of SNARE(Ernst and Brunger, 2003) and Gβ_1_γ_2_ (Sondek et al., 1996a) have been determined alone and with other protein binding partners (Chen et al., 2002; Huang et al., 2019; Lambright et al., 1996; Maeda et al., 2018), and have not exhibited large conformational changes. Therefore the crystal structures of SNARE and Gβ_1_γ_2_ using the ROSETTADOCK (Chaudhury et al., 2011; Gray et al., 2003) protocol of ROSETTA (Bender et al., 2016), as guided by the interaction sites identified by peptide array analysis and mutagenesis. Two regions of the Gβ_1_γ_2_ were probed - the coiled-coil region and the propeller region on the far side of Gβ_1_ away from the coiled-coil. These regions were docked with the C-terminus of SNAP-25 and near residues 151-164 of SNAP-25, both found to be sensitive to mutation for complex formation.

The generated models were evaluated by the total energy of the complex and interface energy between the two binding partners (**Fig 5**). The lowest interface energy models were used as a point of reference and the root mean square deviation (RMSD) was calculated to these models (**Fig 5a**) to determine the convergence of the independent docking runs.

**Fig 5.**
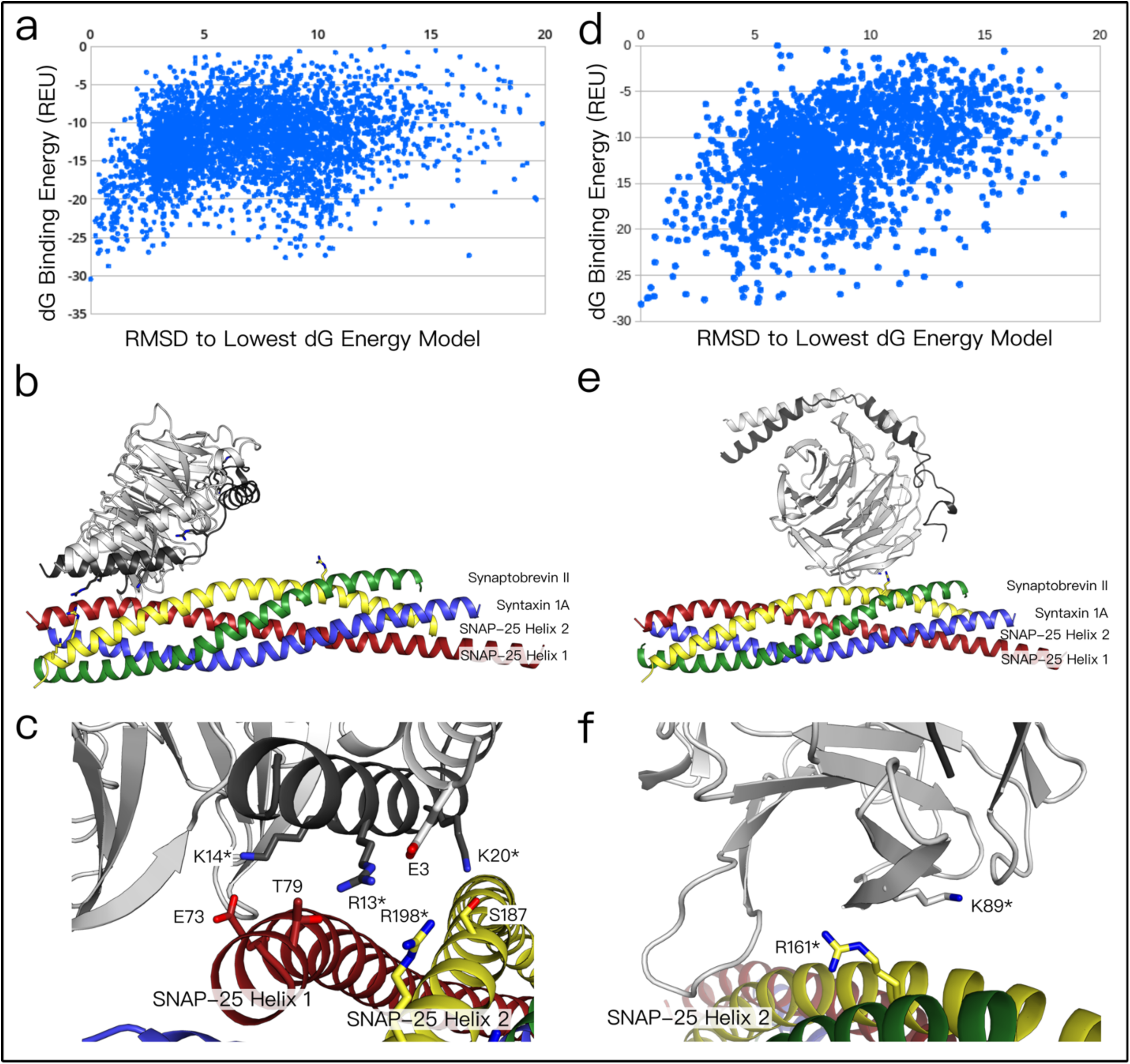
Computational docking of Gβγ to SNARE. **A)** Plot of the ΔG binding energy versus Cα RMSD to lowest energy model of SNARE/Gβγ complex at site 1. The structure of Gβγ (pdbid: 6crk) was computationally docked to the structure of SNARE (pdbid:1n7s) using RosettaDock. The lowest energy model, by binding energy, is shown in (B). **B)** Gβγ/SNARE complex lowest energy model at site 1, by dG binding energy. Complex colored as: Gβ - white, Gγ -black, green – Synaptobrevin II, blue – Syntaxin 1A, red – SNAP-25 (helix 1), yellow – SNAP-25 (helix 2). Residues shown as sticks in the model were shown in to be sensitive to mutagenesis. **C)** Interface of model in (B) - the lowest energy model places the residues shown to be critical on Gγ (R13, K14, K20 and R27) by the interface of the interaction. R198, on helix 2 of SNAP-25, shown to be sensitive to mutagenesis (starred in fig). **D)** Plot of the ΔG binding energy versus Cα RMSD to lowest energy model of SNARE/Gβγ complex at site 2. **E)** Gβγ/SNARE complex lowest energy model at site 2, same coloring as (B). Residues shown as sticks in the model were shown in to be sensitive to mutagenesis. **F)** Interface of model in (E) - the lowest energy model places a residue shown to be critical on Gβ, K89, by the interface of the interaction. R161, on helix 2 of SNAP-25, shown to be sensitive to mutagenesis (starred in fig).

These docking studies identify two distinct docking modes with good convergence. In the first binding mode, the N-terminus of Gγ binds between the two chains of SNAP25 at the C-terminus of the SNARE complex (**Fig 5b**). This model is consistent with the mutagenesis and prior screens. Our previous work (Zurawski et al., 2016) identified SNAP-25^R198^ as a critical residue for the formation of the Gβγ-SNARE complex. In the computational docking model, SNAP-25^R198^ forms a salt bridge with Gβ^E3^. In addition, the peptide mutagenesis of the Gγ subunit suggested that the Gγ residues R13/K14/K20 impact the interaction with SNARE, and these are all located at the proposed interface in the docking model (**Fig 5c**). This model predicts the parallel (as opposed to head-to-tail) orientation of the interactions, with the helical region of Gγ lying flush with the SNARE coiled-coil. The computational model allows for complexin to bind SNARE and Gβγ in a tripartite structure as in the SNARE-complexin structure (Chen et al., 2002).

A second, distinct, docking binding mode that is also consistent with the peptide array analysis and mutagenesis screens, places the Gβγ near the middle of the SNARE complex. In this model, the Gβ^82-96^ and Gβ^124-138^ of the propeller region were located adjacent to SNAP25 residues 151-164, and the critical residue R161. The models with the best convergence place SNAP-25 R161 adjacent to Gβ K89, which was sensitive to alanine mutagenesis. While the residue side chains do not interact in either of the lowest energy models, the positively charged arginine side chain interacts with the backbone of residue 89 or 90.

## Discussion

In general, Gγ subunits may be classified as Gγ_1_-like or Gγ_2_-like. The first category consists exclusively of Gγ_1_ and Gγ_11_, while the ten other Gγ subunits comprise the second category. A critical point of differentiation is found in the CaaX-box motif in each protein: the residue X is a serine in Gγ_1_-like subunits, enabling farnesyltransferase to farnesylate the C-terminus. Gγ_2_-like subunits instead have a leucine residue in this position, conferring geranylgeranylation from geranylgeranyltransferase I instead. It has been previously determined that Gγ_2_-containing Gβγ subunits have substantially higher affinity for the SNARE complex and ability to inhibit exocytosis than Gγ_1_-containing Gβγ subunits (Blackmer et al, 2005, Yoon et al, 2007, Zurawski et al, 2017). Here, we derive new mechanistic insights into this difference in potency. First, we recapitulate the results from our previous peptide inhibition study in which peptides derived from the N-terminus of Gγ_2_ potently inhibit the binding of Gβ_1_γ_2_ to monomeric SNAP25, whereas those derived from the N-terminus of Gγ_1_ do not. We expand upon those studies with a range of potencies for peptides corresponding to different primary sequences within the family of Gγ_2_-like subunits, with N-terminal sequences within Gγ_3 and_ Gγ_12_ providing minimal inhibition, while Gγ_2_, Gγ_8,_ and Gγ_10_ are six to sixteen-fold more potent at inhibiting the binding of Gβ_1_γ_2_ to monomeric SNAP25. From this, we conclude that the primary sequence of the Gγ subunit is contributory towards the binding interface of Gβγ with SNARE, rejecting the notion that the identity of the prenyl group attached to the Gγ is the sole determinant of potency, perhaps due to more efficient membrane targeting.

Scanning alanine mutagenesis of the Gγ_2_ N-terminal peptide suggests that residue R13 and K14 are critical for binding to t-SNARE, as the interaction is abolished when these residues are mutated to Ala (**Fig. 3**). Alignment of primary sequences within the group of Gγ_2_-like subunits using Clustal Omega shows a high degree of conservation of these key residues (**Fig. 6**) between each Gγ subunit. Intriguingly, the most potent N-terminal peptide derived from the primary sequence of Gγ_10,_ was the least conserved by far, featuring a weakly negatively-charged Gln instead of the positively-charged R and K residues. However, this may be consistent with previously published binding data, in which seven of nine Gβγ-binding residues on SNAP25 are R or K residues (Wells et al, 2012). This may imply that Gγ_10_ interacts with SNARE via structurally unique residues compared to other Gβγ heterodimers, since its interaction is predicted from these studies.

**Fig 6.**
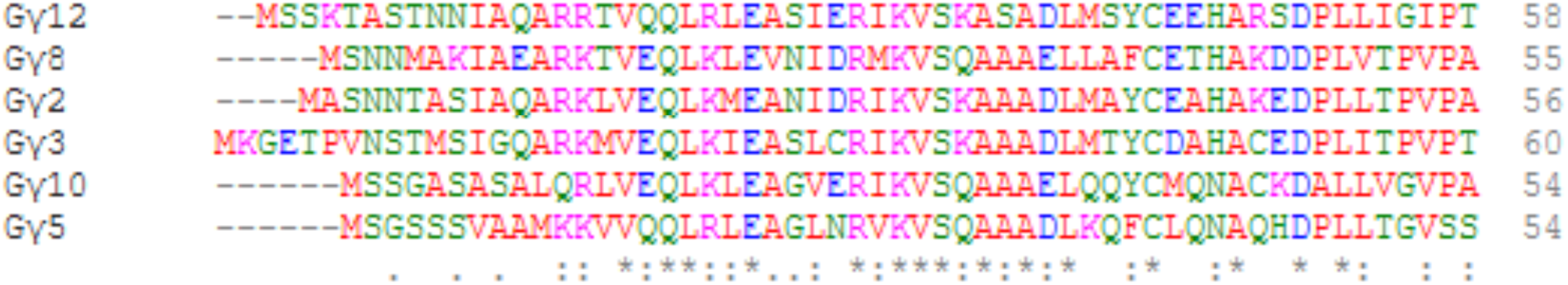
Primary sequence alignment of Gγ_2_-like subunits.

This study, and previous studies produced by our group, show an importance of positively-charged basic arginine and lysine residues on both Gγ and SNAP25 (Wells et al, 2012). Despite this, a predominately positively-charged binding surface of both effectors would reasonably be anticipated to be repulsive. The identity of the negatively charged residues required for the Gβγ-SNARE interaction is less well-understood. Two such Gβγ-binding residues on SNAP25 are E62, located within the first of two helices SNAP25 contributes to the four-helix bundle, and D99, located within the flexible linker region connecting the first and second helices (Wells et al, 2012). The lack of secondary structure in the region where D99 is located would enable the residue to interface with a series of positively-charged residues identified in this data, including Gγ_2_ R13 and K14. However, since the positively charged K102 on SNAP25 is also a key binding residue in close proximity to D99, it would be predicted that a negatively charged residue would have to be in close proximity to R13 and K14, under this hypothesis. Three such residues exist in close proximity on the Gα-binding surface of Gβ, including D163, D186, and D205. On Gγ_2_, residue E17 is also in close proximity, but the data contradicts this, as mutagenesis of E17 to Ala does not interfere with the binding of Gγ_2_ 13-27 to t-SNARE. The limitations of the ResPep technique, including the limited secondary structure of the 15-mer peptides, render it unable to identify every residue of importance within a given protein-protein interaction.

Given this detailed information on residues on both Gβγ and SNAP25, the ROSETTA modeling provides an intriguing and testable hypothesis for the interaction. The two models are distinct, with different regions of the protein interacting for both components of the full complex. The first model exhibits a 1:1 stoichiometry with a parallel coiled-coil orientation. The model highlights the importance of the C-terminus of SNAP25, a critical binding region for the interaction (Gerachshenko et al, 2005, Yoon et al, 2007, Wells et al, 2012, Zurawski et al, 2016), as well as the N-terminus of Gγ (Fig. 2, Fig. 3), Zurawski et al, 2017), all of which were shown to be important in binding and functional studies. Critically, the model emphasizes a 1:1 stoichiometry, while binding studies between labeled Gβ_1_γ_1_ and SNAP25 support two sites of interaction, with one closer to the N-terminus of the second helix of SNAP25 and the other at the C-terminus. Mutation of C-terminal binding residues reduces, but does not abolish the interaction (Wells et al, 2012).

The second model places the propeller domain of Gβ adjacent to the central region of the second SNAP-25 helix. Superposition of the model atop the synaptotagmin-1:ternary SNARE crystal structure (Nature. 2015 Sep 3;525(7567):62-7) demonstrates a steric clash at the primary interface, consistent with a model of orthosteric competition between Gβγ and synaptotagmin 1 C2AB for binding to this region predicted from biochemical experiments (Blackmer et al, 2005, Yoon et al, 2007, Zurawski et al, 2017, Zurawski et al., 2019). The Gβγ-SNARE interaction is blocked in cellular studies through the use of C-terminal peptides derived from GRK2 (Blackmer et al, 2001), for which a crystal structure of the full GRK2 in complex with Gβγ exists (Mol Pharmacol. 2011 Aug;80(2):294-303). While the binding region of GRK2 is not the same as the second model, steric overlap of ternary SNARE and the GRK2 peptide occurs. Neither model features any sites of interaction adjacent to residue W332 of Gβ, but binding studies show that the W332A mutation significantly increases the affinity of Gβγ for t-SNARE complexes consisting of syntaxin1A and SNAP25 (Zurawski et al, 2017), however this may be allosteric in nature.

## Conclusions

The interaction of the G-protein βγ dimer with the SNARE complex is an important regulatory mechanism inhibiting exocytosis. In this study, we probed the regions of Gβγ important for its binding to the SNARE complex. Systematic peptide screening revealed two regions, consistent across the family of five Gβ and twelve Gγ proteins, that can bind to tSNARE. These two regions, the Gβγ coiled-coil domain and the Gβ propeller region, are spatially separated and are unable to act upon the same SNARE. Alanine scanning found that positively charged residues, on both Gβ and Gγ, are important for governing this interaction. Computational docking allowed for the systematic peptide analysis to be evaluated in scope of the entire complex. Analysis of the modeling was in agreement with the peptide screening, there are two distinct binding modes, using two different regions on both Gβγ and SNARE. This may allow for the formation of a higher order complex in vivo. These data, and the models based on them, clearly provide a roadmap for further experiments probing the nature of the interface. Obviously, it would be of great interest to determine the crystal structure to gain a high-resolution view of the Gβγ-SNARE interaction.

## Materials and Methods

### Materials

Primary antibodies SNAP25 (SC-376713) was from Santa Cruz Biotech. HRP-conjugated secondary antibodies (anti-rabbit: 5520-0337, anti-mouse: 55220-0341) were from Seracare (Milford, MA). Enhanced chemiluminescence substrate (34580) was from ThermoFisher (Waltham, MA).

### tSNARE and SNAP25 expression and purification

tSNARE was expressed and purified as described (Zurawski et al., 2017a).

### Peptide Array analysis and Far-western blot

Peptide array synthesis was performed using the ResPep SL peptide synthesizer (Intavis AG, Koeln, Germany) according to standard SPOT synthesis protocols (Frank, 2002; Yim et al., 2015b). Indicated protein peptides containing 15 amino acids were directly coupled to membranes via the C-terminus during synthesis. Dried membranes with peptides were soaked in 100% ethanol for 5 min then rehydrated in water (twice 5 min). The membranes were blocked for 1 h in Tris-buffered saline (TBS) with 5% milk and 0.05% Tween 20 (Sigma-Aldrich, St. Louis, MO) and washed 3 times (5 min each) in TBS with 0.05% Tween 20 (TBS-T). The membranes were incubated overnight at 4°C with t-snare protein at a final concentration of 0.5 µM in 20 mM MOPS, pH 7.5 buffer containing 150 mM NaCl and 2 mM TCEP. The next morning, membranes were washed 3 times (5 min each) in TBS-T buffer and incubated with primary antibody (SNAP-25) at 1:1,000 dilution in TBS-T for 1 h. Spots were detected using HRP-conjugated secondary antibodies as described by the manufacturer. For densitometric analysis, the signal in individual dots was quantified as a percentage of total density detected on the membrane (Quantity One and Image Lab, Bio Rad (Hercules, CA), Image J (Schneider et al., 2012)). This allowed for comparison of intensities across all peptide experiments. Statistical analysis was performed using one-way ANOVA followed by Dunnett’s post-hoc test using GraphPad Prism 8.0.

### Alphascreen competition-binding assays

Alphascreen luminescence measurements were performed as described (Zurawski et al., 2017a). Briefly, biotinylated SNAP-25 was diluted to a 5 concentration of 100 nM in assay buffer (20 mM HEPES, pH 7.0, 10 mM NaCl, 40 mM KCl, 5% glycerol, and 0.01% Triton X-100). An EC80 concentration of 180 nM purified His6-Gβ_1_γ_2_ was made in assay buffer. Peptide stocks in DMSO were spotted onto 384-well white OptiPlates (PerkinElmer Life Sciences) at concentration ranges of 1 nM to 100 M using a Labcyte Echo 555 Omics acoustic liquid handler (Labcyte, Sunnyvale, CA), with DMSO being back-added to a final concentration of 0.1%. 4 ul of Gβ_1_γ_2_ solution was incubated with peptide for 5 min while shaking. After 5 min, 1 ul of biotinylated SNAP25 was added to a final concentration of 20 nM. Subsequent to incubation while shaking for an additional 5 min, 10 ul of Alphascreen Histidine Detection Kit (nickel chelate) acceptor beads were added to a final concentration of 20 ug/ml in assay buffer. The assay plate was agitated in dim green light for 30 min. At that point, Alphascreen Streptavidin Donor Beads were added to a final concentration of 20 ug/ml in dim green light. All aqueous solutions in this assay were manipulated by a Velocity 11 Bravo liquid handler (Agilent Automation Solutions, Santa Clara, CA). The final volume in the assay plate was 25 ul. After being spun down briefly to settle all fluid at the bottom of the well, plates were incubated for an additional 1 h at 27 °C before being read in the EnSpire. 20 nM biotinylated recombinant glutathione S-transferase in place of SNAP-25 with Gβ_1_γ_2_ were used as a negative control for nonspecific binding in each assay. IC50 concentrations for each peptide were determined by sigmoidal dose-response curve-fitting with variable slope. To have a strong signal in the Alphascreen competition-binding assay that could still be competed with a peptide, we used an EC80 concentration (180 nM) of His6-tagged Gβ_1_γ_2_ combined with 20 nM recombinant human SNAP25 biotinylated on primary amine residues with NHS-biotin and increasing concentrations of peptides corresponding to a region on Gγ_2_ in the Alphascreen bead assay. As a negative control, peptides were tested for their ability to disrupt a second Alphascreen assay in which donor and acceptor beads were reacted with 50 nM concentrations of a peptide consisting of a biotinylation site and a His-tag peptide (PerkinElmer Life Sciences).

### Computational docking

Using ROSETTA we applied the previously published protocols ROSETTADOCK to computationally dock the crystal structures of SNARE and Gβ_1_γ_2_. The experimentally determined SNARE structure was mutated at nine positions, from isoform 2 to isoform 1, to match the peptide screen sequence. Both structures were relaxed independently prior to docking. Rigid-body docking with sidechain flexibility was carried out at four distinct positions: placing the Gβ_1_γ_2_ coiled-coil region parallel and anti-parallel to the C-terminal region of SNAP-25, and placing the Gβ^82-96^ and Gβ^124-138^ of the propeller region located adjacent to SNAP25 residues 151-164 in two distinct orientations, with the Gβ_1_ propeller orientation rotated 180 degrees. Approximately 2000 models were generated for each position. The generated models were evaluated by total complex energy and interface energy between the two binding partners using the REF2015 score function. The interface energy was calculated by determining the difference between the score of the bound complex versus the score of the separated complex with the side chains allowed to repack.

